# MicroRNAs as potential therapeutics to enhance chemosensitivity in advanced prostate cancer

**DOI:** 10.1101/273284

**Authors:** Hui-Ming Lin, Iva Nikolic, Jessica Yang, Lesley Castillo, Niantao Deng, Chia-Ling Chan, Nicole K. Yeung, Eoin Dodson, Benjamin Elsworth, Calan Spielman, Brian Y. Lee, Zoe Boyer, Kaylene J. Simpson, Roger J. Daly, Lisa G. Horvath, Alexander Swarbrick

**Author notes:** **Corresponding authors:** Associate Professor Alexander Swarbrick;Tumour Progression Laboratory, Cancer Division, The Kinghorn Cancer Centre, Garvan Institute of Medical Research, 384 Victoria Street, Darlinghurst, New South Wales 2010, Australia., Professor Lisa G. Horvath; Department of Medical Oncology, Chris O’Brien Lifehouse, PO BOX M33, Missenden Road, Camperdown, New South Wales 2050, Australia.

## Abstract

Docetaxel and cabazitaxel are taxane chemotherapy treatments for metastatic castration-resistant prostate cancer (CRPC). However, therapeutic resistance remains a major issue. MicroRNAs are short non-coding RNAs that can silence multiple genes, regulating several signalling pathways simultaneously. Therefore, synthetic microRNAs may have therapeutic potential in CRPC by regulating genes involved in taxane response and minimise compensatory mechanisms that cause taxane resistance. To identify microRNAs that can improve the efficacy of taxanes in CRPC, we performed a genome-wide screen of 1280 microRNAs in the CRPC cell lines PC3 and DU145 in combination with docetaxel or cabazitaxel treatment. Mimics of miR-217 and miR-181b-5p enhanced apoptosis significantly in PC3 cells in the presence of these taxanes. These mimics downregulated at least a thousand different transcripts, which were enriched for genes with cell proliferation and focal adhesion functions. Individual knockdown of a selection of 46 genes representing these transcripts resulted in toxic or taxane sensitisation effects, indicating that these genes may be mediating the effects of the microRNA mimics. A range of these genes are expressed in CRPC metastases, suggesting that these microRNA mimics may be functional in CRPC. With further development, these microRNA mimics may have therapeutic potential to improve taxane response in CRPC patients.

## Introduction

Prostate cancer is the second most frequently diagnosed cancer in men worldwide, and the third leading cause of male cancer death in developed countries^1^. Despite the rise in new therapeutics for metastatic castration-resistant prostate cancer (CRPC) such as novel anti-androgens, radium-223 and PARP inhibitors^2,3^, the taxanes docetaxel and cabazitaxel are the standard of care chemotherapy treatments for CRPC. For more than a decade, docetaxel has remained as the first line cytotoxic treatment for CRPC^4,5^, and is now increasingly used in the metastatic castration-sensitive setting^6,7^. However, only ∼50% of patients respond to docetaxel, and responders eventually develop resistance^4,5^. Cabazitaxel, a second-generation taxane, improves the survival of patients with docetaxel-resistant CRPC, but was not superior to docetaxel and thus remains as second line treatment with a response rate of ∼60%^8,9^. Overall, new therapeutic strategies are required to overcome taxane resistance and improve patient outcome.

MicroRNAs are short non-coding RNAs (∼22 nucleotides) that regulate gene expression post-transcriptionally by forming an RNA-induced silencing complex which represses translation or degrades messenger RNA (mRNA)^10^. The complex is formed by a microRNA binding to the mRNA at a complementary ‘seed’ sequence on the 3’ untranslated region of the mRNA, together with Argonaute proteins. Binding can occur with imperfect base pairing, thus a single microRNA can negatively regulate hundreds of different genes, and the mRNA of a single gene can be targeted by different microRNAs. Many microRNAs have tissue-specific expression^11^, and over two thousand different microRNAs have been identified in humans^12^.

The discovery of oncogenic and tumour-suppressor microRNAs, and the ability to manipulate cellular microRNA levels with modified oligonucleotides that mimic or inhibit their function has lead to extensive research and development of microRNAs as therapeutics^13,14^. By using a single microRNA to silence multiple genes, several signalling pathways can be regulated simultaneously and may thus minimise compensatory mechanisms that cause therapeutic resistance. Most microRNA-based therapeutics are still in the pre-clinical stages of research^14^. A few have completed Phase 1 or Phase 2 clinical trials with positive results, such as Miravirsen (miR-122 inhibitor) for hepatitis C viral infection^15^ and TagomiR (miR-16 mimic) for malignant pleural mesothelioma^16^.

A potential therapeutic application of microRNAs is to combine microRNA therapy with taxane chemotherapy to overcome chemoresistance. The availability of large libraries of microRNA mimics or inhibitors enables the use of genome-wide screens to identify microRNAs that can increase the sensitivity of cancer cells to a drug. This approach was commonly used with small-interfering RNA (siRNA) libraries to identify synthetic lethal genes^17^ but has been demonstrated with microRNA libraries. For example, Lam *et al* identified 19 microRNAs that sensitised colorectal cancer cells to a BCL2 inhibitor from a screen of 810 microRNA mimics combined with the drug^18^.

To identify microRNAs that have therapeutic potential to overcome resistance to docetaxel and cabazitaxel in CRPC, we performed a genome-wide functional screen of CRPC cell lines PC3 and DU145, with microRNA mimics or inhibitors of 1280 microRNAs in combination with docetaxel or cabazitaxel treatment. MicroRNA mimics act as mature microRNAs, whereas microRNA inhibitors block the activity of endogenous microRNAs within a cell. The screen and the effects of the mimics/inhibitors on the cells without taxane treatment are described by Nikolic *et al* 2017^19^. Here, we present the findings from combining the screen with taxane treatment, where the effects and gene targets of two hits from the screen were further characterised.

## Results

### Identification of microRNAs that are taxane sensitisers

A high-throughput functional screen testing microRNA mimics or inhibitors of 1280 microRNAs in PC3 and DU145 cancer cell lines was performed to identify microRNA mimics or inhibitors that can increase the sensitivity of the cells to docetaxel or cabazitaxel. Non-targeting siRNA was used as the control for the microRNA mimics or inhibitors. The cells in the screen were treated with sub-optimal doses of docetaxel and cabazitaxel equivalent to an IC20 dose (drug concentration where response is reduced by 20%). Given that these doses of taxanes decreased cell viability by 20% in non-targeting control cells, we considered a microRNA mimic or inhibitor as a taxane sensitiser if it decreased cell viability by more than 20% with taxane treatment – i.e. viability of taxane-treated cells transfected with mimic or inhibitor <80% of the viability of taxane-treated cells transfected with non-targeting control, and viability of vehicle-treated cells transfected with mimic or inhibitor >80% of the viability of vehicle-treated cells transfected with non-targeting control.

These thresholds produced many hits among the microRNA mimics – 102 docetaxel sensitisers and 112 cabazitaxel sensitisers for PC3 cells, 61 docetaxel sensitisers and 49 cabazitaxel sensitisers for DU145 cells (Figure 1a). Most of these hits only caused a modest effect with taxane treatment (70-80% viability), and only four reduced cell viability by more than 50% (Figure 1a). None of the hits caused the same level of toxicity, with or without taxane treatment, as that of the positive control, PLK1 (polo-like kinase 1) siRNA, where the viability of PC3 and DU145 cells transfected with PLK1 siRNA was 22% and 6% respectively.

**Figure 1.**
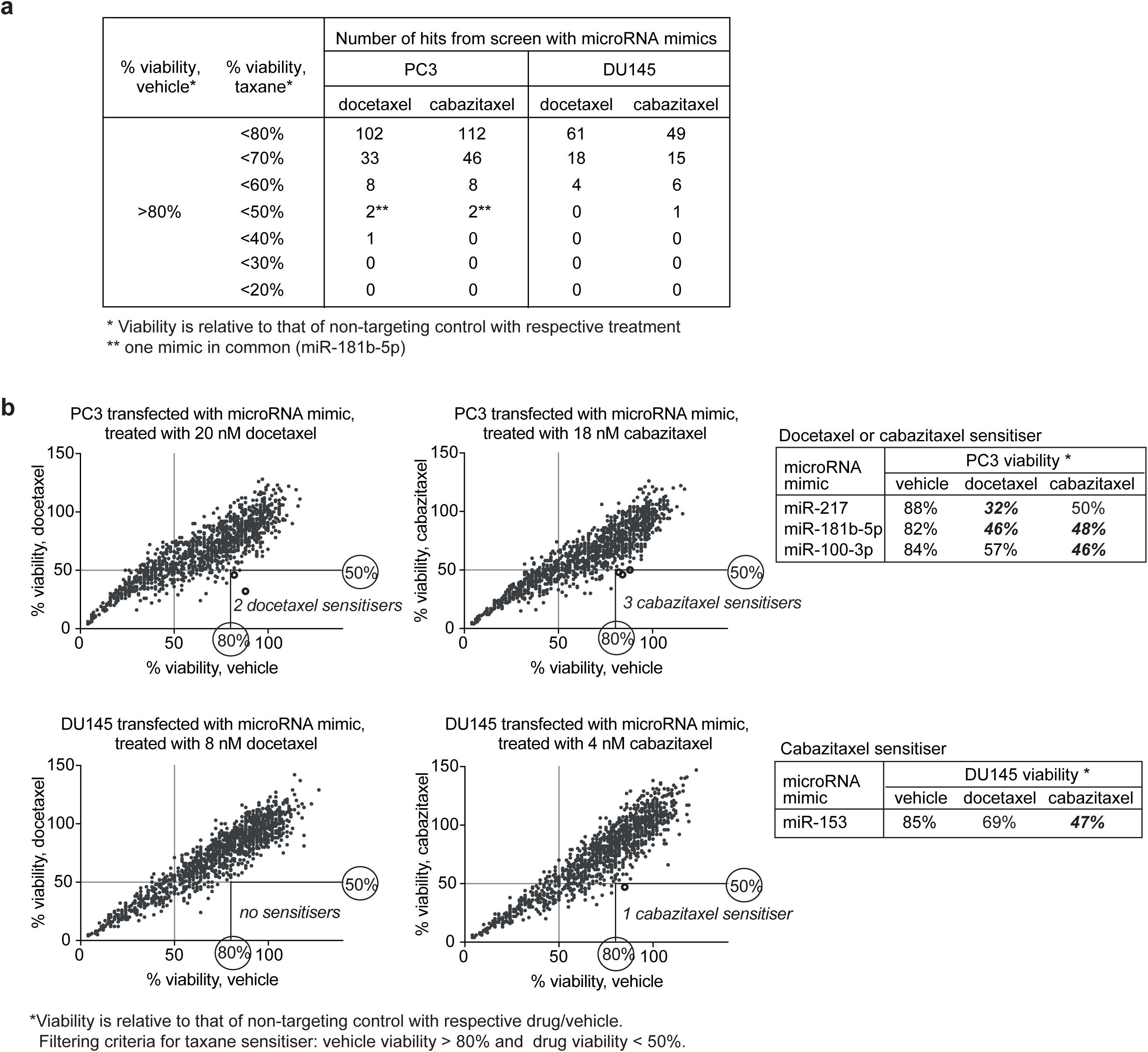
Identification of microRNA mimics with taxane sensitisation effects in PC3 and DU145 cell lines in a functional screen testing 1280 microRNA mimics individually in combination with docetaxel or cabazitaxel treatment: (a) Number of hits at different thresholds of taxane sensitisation; (b) Viability plots showing the thresholds for identifying the strongest taxane sensitisers in PC3 and DU145. *Viability is relative to that of non-targeting control with respective drug/vehicle. Filtering criteria for taxane sensitiser: vehicle viability > 80% and drug viability < 50%.

The number of microRNA inhibitors with taxane sensitisation effects were far less compared to the microRNA mimics – 71 docetaxel sensitisers and 37 cabazitaxel sensitisers for PC3 cells, 29 docetaxel sensitisers and 22 cabazitaxel sensitisers for DU145 cells (Supplementary Figure S1A). Unlike the microRNA mimics, none of the microRNA inhibitors reduced cell viability by more than 50% (Supplementary Figure S1B). The strongest taxane sensitisation effect by any of the inhibitors was 40% reduction in cell viability with cabazitaxel treatment, and 30% reduction in cell viability with docetaxel treatment.

The microRNA mimics with the strongest taxane sensitising effects – causing more than 50% cell death, were selected for further investigation. These consisted of miR-217, miR-181b-5p, miR-100-3p and miR-153 mimics (Figure 1b). MiR-217, miR-181b-5p, and miR-100-3p increased the sensitivity of PC3 cells to either or both taxanes, whereas miR-153 increased the sensitivity of DU145 to cabazitaxel in the screen.

### Effect of microRNA mimics on cell viability

Further validation with a range of taxane doses to generate dose response curves showed that miR-217 and miR-181b-5p mimics decreased the IC50 (drug concentration where response is reduced by 50%) of docetaxel or cabazitaxel, or enhanced the maximum response to the taxanes (lowest cell viability among the drug concentrations) (Figure 2a). These mimics also modestly reduced cell number in the absence of taxanes, as cells transfected with the mimics grew more slowly than those transfected with the non-targeting control (Figure 2b).

**Figure 2.**
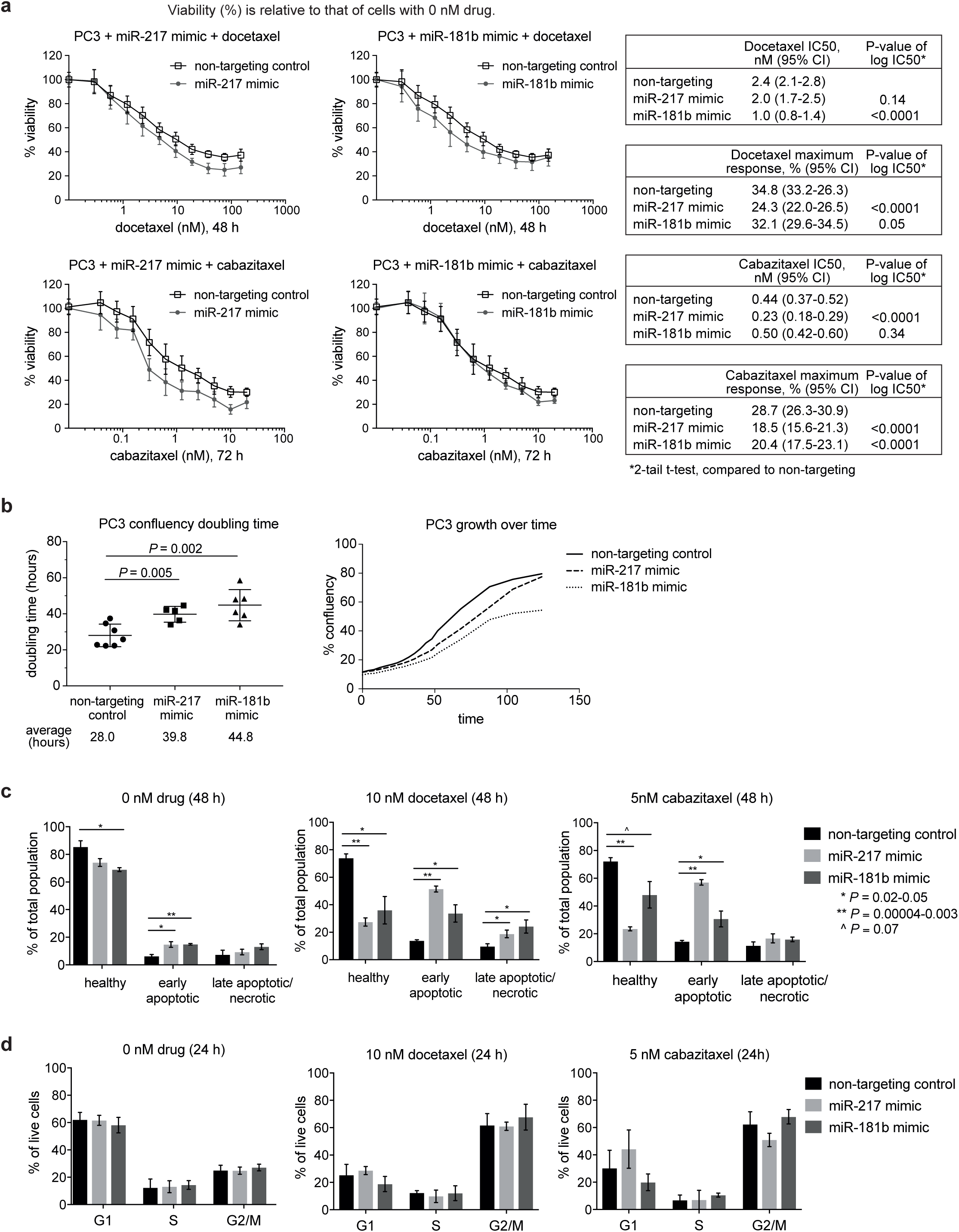
Assessment of cell viability, apoptosis, and cell cycle phases in PC3 cells transfected with miR-217 or miR-181b-5p mimic: (a) Docetaxel and cabazitaxel dose response curves (mean ± standard deviations of 3 experiments with 4 replicate samples each); (b) Doubling times measured using Incucyte Zoom, and an example of growth curves from one experiment; (c) Percentage of apoptotic cells, quantitated by flow cytometry with annexin-V-FITC and propidium iodide labelling (mean ± standard deviations of 3 experiments); (d) Percentage of cells in cell cycle phases, quantitated by flow cytometry with propidium iodide staining (mean ± standard deviations of 3 experiments).

MiR-100-3p and miR-153 mimics also reduced cell number in transfected PC3 and DU145 cells in the absence of taxanes, but did not alter the IC50 or the maximum response of cabazitaxel in PC3 and DU145 respectively (Supplementary Figure S2). Therefore, miR-217 and miR-181b-5p were selected for further study as potential therapeutics.

The slower growth rate of PC3 cells when transfected with the mimics may be partly attributed to enhanced apoptosis, as the percentage of apoptotic cells was significantly higher among cells transfected with the mimics compared to those transfected with the non-targeting control (Figure 2c – 0 nM drug, P<0.05). Addition of docetaxel or cabazitaxel significantly increased apoptosis in cells transfected with the mimics, compared to those transfected with non-targeting control (Figure 2c – 10 nM docetaxel and 5nM cabazitaxel, P<0.05).

There were no significant differences in the percentage of cells in the cell cycle phases of G1, S, and G2-M for cells transfected with the mimics compared to those transfected with the non-targeting control, with or without taxane treatment (Figure 2d). Docetaxel and cabazitaxel caused cell cycle arrest in the G2-M phase as expected, without any significant differences in cell numbers between mimics and non-targeting control (Figure 2d, P>0.05). These findings indicate that enhanced apoptosis by the mimics occurred through mechanisms that did not involve alterations to the taxanes’ induction of G2-M cell cycle arrest.

The effects of miR-217 and miR181b-5p on cell viability were only specific to PC3 and not DU145, as these mimics did not alter the taxane dose response curves or enhanced apoptosis in DU145 cells (Supplementary Figure S3).

### Genes regulated by microRNA mimics

To identify genes and pathways that were regulated by the microRNA mimics, RNA-sequencing was performed on PC3 cells transfected with miR-217 or miR-181b-5p mimics. Filtered reads were mapped to 27,718 genes. The number of transcripts significantly downregulated in cells transfected with miR-217 or miR-181b-5p mimics compared to non-targeting control were 1295 and 1355 respectively (adjusted P<0.05), of which 402 (∼30%) were in common. According to the microRNA target database TarBase (version 7)^20^, 48 (4%) and 258 (19%) of the transcripts downregulated by miR-217 and miR-181b-5p mimics respectively were experimentally-supported targets. Our own database^19^ predicted that 490 (38%) and 518 (38%) of the transcripts downregulated by miR-217 and miR-181b-5p mimics respectively were direct targets of these mimics. The target predictions made by our database were the combined results from established target prediction algorithms by StarBase, MIRTarBase, TargetScan, DIANA and MIRDB^19^.

The Molecular Signatures Database^21^ was used to identify biological themes that were enriched among the downregulated transcripts, as these themes may provide insights into the function of the gene targets and hence the mechanism of the microRNA mimics. Several of the biological themes that were significantly enriched among the downregulated transcripts in cells transfected with either miR-217 or miR-181b-5p mimics were related to cell proliferation and focal adhesion (Table 1).

**Table 1.** Biological themes enriched among RNA transcripts downregulated by microRNA mimics, based on significant overlaps with gene sets from the Molecular Signatures Database (MSigDB).

| **miR-217 mimic-transfected PC3 cells (1274 genes recognised by MSigDB)** |  |  |  |
| --- | --- | --- | --- |
| *Top 10 Hallmark gene sets* | *Genes in gene set* | *Genes in overlap* | *P-value* |
| 1 **E2F targets**<sup>a</sup> | 200 | 64 | 2.2x10<sup>-49</sup> |
| 2 **G2M checkpoint**<sup>a</sup> | 200 | 61 | 1.0x10<sup>-45</sup> |
| 3 **Mitotic spindle**<sup>a</sup> | 200 | 44 | 1.5x10<sup>-25</sup> |
| 4 **Epithelial-mesenchymal-transition**<sup>b</sup> | 200 | 40 | 8.3x10<sup>-23</sup> |
| 5 MYC targets | 200 | 38 | 6.5x10<sup>-21</sup> |
| 6 Estrogen response late | 200 | 29 | 3.9x10<sup>-13</sup> |
| 7 DNA repair | 150 | 25 | 6.7x10<sup>-13</sup> |
| 8 Apical junction | 200 | 27 | 1.4x10<sup>-11</sup> |
| 9 Myogenesis | 200 | 24 | 2.1x10<sup>-9</sup> |
| 10 UV response | 144 | 20 | 3.6x10<sup>-9</sup> |
| *Top 10 KEGG gene sets* | *Genes in gene set* | *Genes in overlap* | *P-value* |
| 1 **DNA replication**<sup>a</sup> | 36 | 15 | 1.3x10<sup>-14</sup> |
| 2 **Cell cycle**<sup>a</sup> | 128 | 24 | 1.3x10<sup>-13</sup> |
| 3 **Focal Adhesion**<sup>b</sup> | 201 | 28 | 2.7x10<sup>-12</sup> |
| 4 Nucleotide excision repair | 44 | 13 | 1.3x10<sup>-10</sup> |
| 5 **ECM receptor interaction**<sup>b</sup> | 84 | 16 | 1.2x10<sup>-9</sup> |
| 6 Pathways in cancer | 328 | 31 | 3.9x10<sup>-9</sup> |
| 7 Mismatch repair | 23 | 9 | 5.4x10<sup>-9</sup> |
| 8 Pathogenic E. Coli infection | 59 | 13 | 6.7x10<sup>-9</sup> |
| 9 Ubiquitin mediated proteolysis | 138 | 19 | 1.0x10<sup>-8</sup> |
| 10 Oocyte meiosis | 114 | 17 | 1.8x10<sup>-8</sup> |
| **miR-181b-5p mimic-transfected PC3 cells (1337 genes recognised by MSigDB)** |  |  |  |
| *Top 10 Hallmark gene sets* | *Genes in gene set* | *Genes in overlap* | *P-value* |
| 1 MYC targets | 200 | 105 | 4.6x10<sup>-106</sup> |
| 2 **G2M checkpoint**<sup>a</sup> | 200 | 76 | 2.0x10<sup>-63</sup> |
| 3 **E2F targets**<sup>a</sup> | 200 | 72 | 3.6x10<sup>-58</sup> |
| 4 **Mitotic spindle**<sup>a</sup> | 200 | 56 | 1.1x10<sup>-38</sup> |
| 5 MTORC1 signaling | 200 | 56 | 1.1x10<sup>-38</sup> |
| 6 Oxidative phosphorylation | 200 | 49 | 5.0x10<sup>-31</sup> |
| 7 **Epithelial mesenchymal transition**<sup>b</sup> | 200 | 46 | 6.4x10<sup>-28</sup> |
| 8 Fatty acid metabolism | 158 | 35 | 5.2x10<sup>-21</sup> |
| 9 Glycolysis | 200 | 37 | 2.7x10<sup>-19</sup> |
| 10 Adipogenesis | 200 | 31 | 3.4x10<sup>-14</sup> |
| *Top 10 KEGG gene sets* | *Genes in gene set* | *Genes in overlap* | *P-value* |
| 1 Ribosome | 88 | 36 | 4.5x10<sup>-32</sup> |
| 2 Huntingtons disease | 185 | 38 | 1.9x10<sup>-21</sup> |
| 3 Oocyte meiosis | 114 | 30 | 1.7x10<sup>-20</sup> |
| 4 Spliceosome | 128 | 31 | 5.5x10<sup>-20</sup> |
| 5 **Cell cycle**<sup>a</sup> | 128 | 30 | 6.0x10<sup>-19</sup> |
| 6 Parkinsons disease | 133 | 29 | 1.9x10<sup>-17</sup> |
| 7 Alzheimers disease | 169 | 32 | 3.4x10<sup>-17</sup> |
| 8 **Focal adhesion**<sup>b</sup> | 201 | 34 | 1.3x10<sup>-16</sup> |
| 9 Pathways in cancer | 328 | 40 | 2.8x10<sup>-14</sup> |
| 10 Oxidative phosphorylation | 135 | 25 | 1.6x10<sup>-13</sup> |

<sup>a</sup> cell proliferation theme
<sup>b</sup> focal adhesion theme

To determine if the downregulated transcripts mediate the taxane sensitisation effect of the microRNA mimics, siRNA SMARTpool knockdown of 51 genes individually in PC3 cells in combination with docetaxel or cabazitaxel treatment was performed with miR-217 and miR-181b mimics as positive controls (full names listed in Supplementary Table S1). These 51 genes were selected on the basis of their association with the enriched biological themes, upregulation of phosphorylation/protein levels in a docetaxel-resistant PC3 model compared to docetaxel-sensitive PC3 (Supplementary Table S2)^22^, or significant downregulation of transcripts by the mimics. Interestingly, one of these genes is PLK1, whose knockdown is known to be lethal and hence its siRNA is commonly used as a positive control of transfection and in our microRNA screen.

SiRNA knockdown of four genes resulted in significant toxicity – PLK1, FBXO5 (F-box protein 5), TUBA1A (tubulin alpha 1a), RPLP0 (ribosomal protein lateral stalk subunit P0) (Figure 3). The viability of cells with knockdown of either of these four genes was 20-40% of that of control (non-targeting siRNA) without taxane treatment, and taxane treatment did not decrease their viability further (Figure 3).

**Figure 3.**
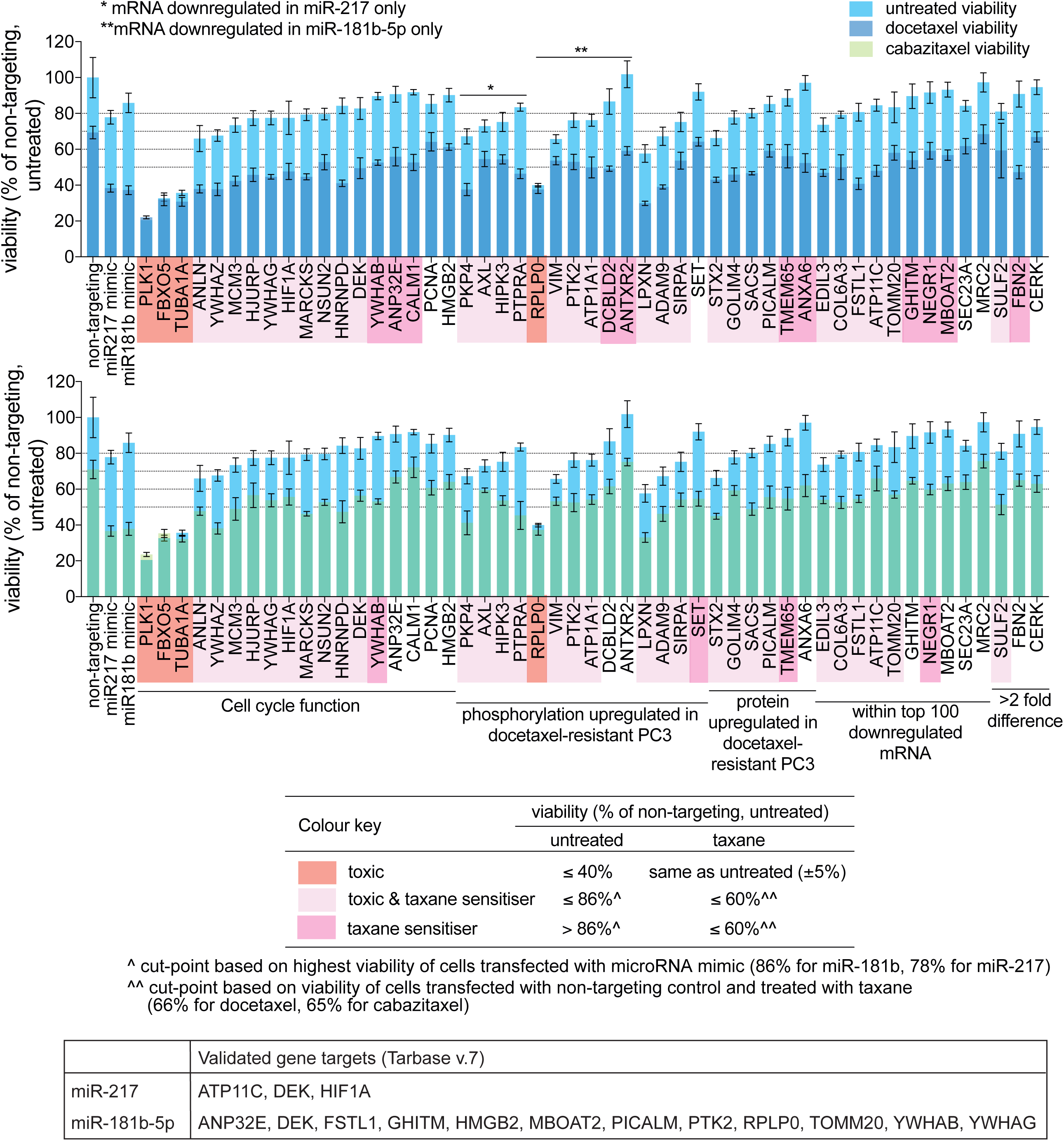
Effects of siRNA knockdown of 51 gene targets of miR-217 or miR-181b-5p mimics on the docetaxel or cabazitaxel sensitivity of PC3 cells. Cells were treated with 10nM docetaxel or 5nM cabazitaxel after siRNA transfection. Barplots show viability of siRNA-transfected cells, with or without taxane treatment, relative to that of cells transfected with non-targeting siRNA control without taxane treatment. Datapoints are mean ± standard deviations of 4 replicate samples.

Twelve genes were taxane sensitisers upon siRNA knockdown, where their knockdown increased docetaxel or cabazitaxel sensitivity without an effect in the absence of the taxanes – e.g. CALM1 (calmodulin 1), TMEM65 (transmembrane protein 65), FBN2 (fibrillin 2) (Figure 3). However, the taxane sensitisation effects exerted by the knockdown of these 12 genes were weaker than that of miR-217 and miR-181b-5p mimics. When compared to taxane-treated control, taxane-treated cells with knockdown of either of these 12 genes had 10-20% lower viability whereas taxane-treated mimic-transfected cells had ∼30% lower viability (Figure 3).

Thirty genes were both toxic and taxane sensitisers upon siRNA knockdown – e.g. ANLN (anillin actin binding protein), PKP4 (plakophilin 4), STX2 (synthaxin 2) (Figure 3). The knockdown of either of these 30 genes caused various degrees of toxicity which was enhanced by docetaxel or cabazitaxel treatment (Figure 3).

Overall, a total of 46 genes have toxic (4 genes), taxane sensitisation (12 genes) or both effects (30 genes) when silenced by siRNA knockdown. These 46 genes may be direct or indirect targets of the microRNA mimics that contribute to the taxane sensitisation effect of the mimics. The genes that display toxic effects with or without taxane sensitisation upon siRNA knockdown may be mediating the killing effects of the microRNA mimics in the absence of taxane treatment. Of these 46 genes, 13 were experimentally validated targets of miR-217 or miR-181b-5p according to TarBase version 7 (Figure 3).

### Evaluation of gene targets and microRNAs in genomic tumour datasets

To determine if the genes downregulated by miR-217 and miR-181b-5p mimics in PC3 cells are expressed in human CRPC tumours, the mRNA levels of these genes were examined in three different genomic datasets of CRPC tumours (Grasso *et al*, Kumar *et al*, Robinson *et al*)^23-25^. The datasets are available from the cBioPortal database^26^ or NCBI GEO repository. The genes downregulated by the microRNA mimics are referred to as gene targets (direct or indirect) henceforth.

The dataset by Grasso *et al* consists of 34 CRPC metastases of soft tissue organs obtained at rapid autopsy, and 28 non-malignant prostate tissue samples (unmatched) taken from the radical prostatectomy of patients with prostate cancer^23^. Microarray expression data for ∼92% of the gene targets of either miR-217 and miR-181b-5p were available for these tissues, of which 18% of miR-217 gene targets and 13% of miR-181b-5p gene targets were significantly upregulated in metastases compared to normal prostate tissue (adjusted t-test P-value≤0.05, fold≥1.5). Of the 46 gene targets that were shown to have toxic or taxane sensitisation effects upon siRNA knockdown in PC3, ten were upregulated in metastases compared to non-malignant tissues (Figure 4a). The siRNA knockdown of all these ten genes in PC3 enhanced sensitivity to both docetaxel and cabazitaxel (Figure 3). Knockdown of all except SET (SET nuclear proto-oncogene) also resulted in toxicity (Figure 3).

**Figure 4.**
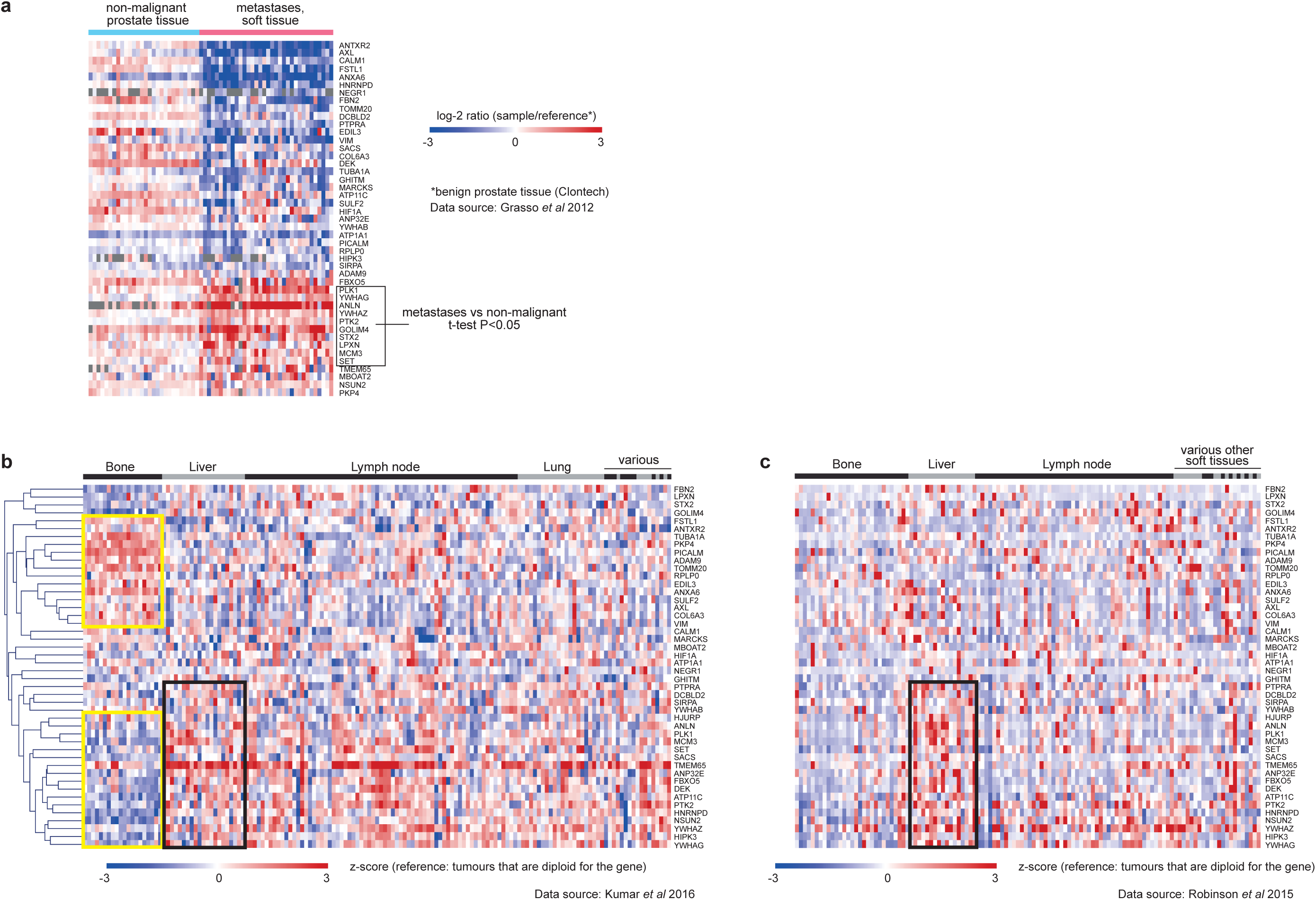
Heatmap of mRNA expression for 46 gene targets of miR-217 or miR-181b-5p mimics in: (a) non-malignant prostate tissue samples from radical prostatectomy of 28 men, and metastatic CRPC samples from rapid autopsy of 34 men; (b) 149 metastatic CRPC samples from rapid autopsy of 63 men; (c) metastatic CRPC biopsies from 118 men. Genes in heatmaps B & C are in the same order.

The dataset by Kumar *et al*^24^ consists of 129 soft tissue and 20 bone CRPC metastases obtained at rapid autopsy. Non-malignant tissue samples were unavailable for comparison. In this dataset, the gene expression profile of the 46 gene targets in bone metastases differed from that of soft tissue metastases (Figure 4b). Distinct clusters of genes with increased or decreased expression were evident among the bone metastases, compared to those derived from soft tissue organs (Figure 4b, outlined in yellow), indicating that the miR-217 and miR-181b-5p mimics may act on different mRNA targets in different metastatic tissues.

Interestingly, the gene expression profile of the 46 gene targets for CRPC metastases in the dataset by Robinson *et al*^25^ differed from that of Kumar *et al* (Figure 4c). The metastases in the dataset by Robinson *et al* consists of 89 soft tissue and 29 bone metastases that were obtained by image-guided biopsy from living individuals at an earlier disease stage of CRPC^25^. The distinct cluster of highly expressed genes observed among the bone metastases in the dataset by Kumar *et al* were not obvious in the dataset by Robinson *et al* (Figure 4c). However, a cluster of upregulated genes in liver metastases could be observed in both datasets (Figure 4b & c, outlined in black). The variation in the mRNA profile in the dataset by Robinson *et al* suggests that the miR-217 and miR-181b-5p mimics will have different mRNA targets at different stages of CRPC.

Overall, these genomic data show that the gene targets of miR-217 and miR-181b-5p mimics are expressed in CRPC metastases, indicating that these microRNA mimics may be functional in CRPC. The microRNA mimics are likely to act on different gene targets according to their expression in the metastatic tissues and disease stage.

## Discussion

Using a high-throughput genome-wide functional screen, we identified two microRNA mimics – miR-217 and miR-181b-5p, which substantially enhance the docetaxel and cabazitaxel sensitivity of the PC3 prostate cancer cell line. An advantage of utilising microRNA mimics as therapeutic agents is their ability to regulate the expression of multiple genes. It is unclear as to how many and which genes are the key direct targets required to elicit the taxane sensitisation effect of the microRNA mimics, as prediction of direct gene targets is difficult due to imperfect base pairing of microRNAs with their target mRNA. Nevertheless, examination of 46 gene targets with toxic or taxane sensitisation effects showed that these genes are expressed in CRPC metastases. Although the expression profile of these genes vary with metastatic site and disease stage, the ability of microRNAs to affect multiple targets and the presence of more than one gene target in CRPC indicates that these microRNA mimics may be effective in all CRPC metastases.

These microRNA mimics appear to enhance the efficacy of the taxanes through mechanisms related to cell proliferation and focal adhesion as these biological themes were enriched among the RNA transcripts downregulated by the microRNA mimics. These mechanisms are consistent with the anti-mitotic and tubulin-targeting action of the taxanes. Taxanes bind directly to microtubules and prevent their disassembly for chromosomal segregation during mitosis, thus resulting in cell cycle arrest in the G2-M phase and induction of apoptosis^27^. Our study showed that knockdown of genes involved in cell proliferation, which are also targets of the microRNA mimics, resulted in taxane sensitisation or toxicity effects. These genes function as cell-cycle related targets of E2F transcription factors, or are involved in the G2-M phase checkpoint or mitotic spindle assembly. Interestingly, some of these genes are known to be over-expressed in cancer or associated with poor prognosis such as PLK1, ANLN and FBXO5^28-30^. Indeed, we observed that both PLK1 and ANLN were significantly upregulated in CRPC metastases compared to non-malignant prostate tissue in the genomic dataset by Grasso *et al*.

Other studies have shown that PLK1 promotes resistance to various chemotherapy agents including docetaxel^31^. Presumably PLK1 caused docetaxel resistance by interfering with docetaxel’s effect on microtubule dynamics, which also affects androgen signalling in prostate cancer^32^. Clinical trials of early PLK1 inhibitors in cancers were disappointing, although the newer inhibitor Volasertib showed encouraging results in combination with cytarabine in leukemia^31^, suggesting that PLK1 inhibitors may be more effective in a combination setting. Indeed, a recent study showed that PLK1 inhibition enhanced the efficacy of the PARP inhibitor, Olaparib, in BRCA-mutant CRPC cell lines and xenografts^33^. Targeting PLK1 via miR-217 or miR-181b mimics is akin to a combination approach as the microRNA mimics target multiple genes, and thus may be more effective than using a single agent PLK1 inhibitor.

Taxanes can also disrupt non-mitotic processes that involve microtubules, such as focal adhesion dynamics^34^. Previously we found that the phosphorylation and protein levels of focal-adhesion and cytoskeletal-related proteins were altered in docetaxel-resistant PC3 and DU145 prostate cancer cells^22^. These alterations appear to contribute to docetaxel resistance, as inhibition of the activity of one of these proteins, PTK2 (protein tyrosine kinase 2, also known as focal adhesion kinase), using tyrosine kinase inhibitors overcame docetaxel resistance and enhanced docetaxel efficacy in xenografts and *ex vivo* cultures of patient-derived prostate tumours^22,35^. In the current study, PTK2 and the other affected focal-adhesion or cytoskeletal-related proteins such as VIM (vimentin) and PKP4 (plakophilin 4) were also targets of the miR-217 or miR-181b-5p mimics, and their knockdown caused toxicity and enhanced taxane sensitivity. Overall, these findings demonstrate that various mechanisms centering on microtubules can influence the activity of taxanes.

The subset of miR-217 and miR-181b-5p gene targets examined in this study display differences in docetaxel and cabazitaxel sensitivity upon siRNA knockdown, supporting the existence of differences in the mechanism of actions between the taxanes despite sharing the same molecular target. Other studies have shown that although both taxanes share similar resistance mechanisms, cabazitaxel also has alternative molecular actions as it is effective in docetaxel-resistant models and an enzalutamide-resistant model that was cross-resistant to docetaxel^36,37^.

To date, there are no studies of miR-217 in prostate cancer. MicroRNA profiling of CRPC showed that miR-217 was not detected in CRPC^38^, and previously we found that miR-217 was not expressed in PC3 cells^*39*^. Studies of various other cancers showed that miR-217 has tumour suppressor properties as miR-217 overexpression inhibited proliferation and invasion, promoted apoptosis, and suppressed xenograft tumour growth^40-44^. Overall, the findings from these studies indicate that miR-217 mimic is ideal as an anti-cancer therapy. Furthermore, some of these studies also found that the levels of miR-217 in tumours were lower than normal tissue, or inversely correlated with tumour stage and survival^42,45,46^. Lower levels of miR-217 in ovarian tumours and leukemia cells was associated with therapeutic resistance^47,48^.

In contrast, results were conflicting from studies on the role of miR-181b-5p in various cancers, with miR-181b-5p displaying oncogenic properties in some studies^49-51^, and acting as a tumour suppressor in others^52-55^. Only one study of miR-181b-5p in prostate cancer has been reported, where suppression of miR-181b expression in PC3 cells induced apoptosis, and inhibited proliferation and invasion^56^. These findings contradict our results, where we showed that miR-181b-5p mimic acted as a tumour suppressor by inhibiting cell growth and enhancing apoptosis.

*In vivo* testing of miR-217 and miR-181b-5p mimics will be required to evaluate their ability to overcome taxane resistance without significant toxicity. The microRNA mimics can be encapsulated in specialised delivery particles labelled with antibodies targeting tumour antigens such as PSMA to enable selective delivery to prostate cancer cells. This approach was demonstrated in the treatment of malignant pleural mesothelioma with TargomiR, which consisted of miR-16 mimics packaged in bacteria-derived nanoparticles labelled with an EGFR-antibody^16^. The Phase 1 study showed that the treatment was well-tolerated and associated with tumour response or disease stabilisation^16^.

In conclusion, we have identified two microRNA mimics that enhanced the efficacy of docetaxel and cabazitaxel in CRPC *in vitro*. With further research and development, these microRNA mimics may have potential to improve taxane chemotherapy response in CRPC patients.

## Methods

### Cell lines, microRNA mimics, and drugs

PC3 and DU145 cell lines were purchased from the American Type Culture Collection (Manassas, VA, USA) and cultured in RPMI 1640 with 10% fetal calf serum, 1.3μM insulin, and 10mM HEPES. These cell lines were authenticated by CellBank Australia using Short Tandem Repeat profiling.

MiR-217 and miR-181b mimics (miRIDIAN microRNA mimics), and ON-TARGETplus non-targeting control siRNA SMARTpool were purchased from GE Healthcare Dharmacon. Docetaxel (clinical-grade) was purchased from Sanofi-Aventis, Australia, and prepared according to manufacturer’s instruction. Cabazitaxel was provided by Sanofi-Aventis and dissolved in 56% ethanol and 10% polysorbate 80 at 5mg/ml. Working stocks of both drugs were prepared in phosphate-buffered saline or culture medium.

### Functional microRNA screen

High-throughput screens of PC3 and DU145 cells with Dharmacon’s miRIDIAN microRNA Mimic and Hairpin Inhibitor libraries (version 16) (GE Healthcare Dharmacon) was performed as described in detail by Nikolic *et al*^19^. Basically, reverse transfection of cells with the libraries was performed in 384-well microplates with DharmaFECT-1 Transfection Reagent. Positive and negative technical controls were included on each library plate, including the ON-TARGETplus non-targeting control siRNA SMARTpool (GE Healthcare Dharmacon) which was used as the control for analysis of the mimics or inhibitors. Culture media was replaced with fresh media at 24 hours post-transfection. Cells were treated with taxanes or vehicle control at 3 days post-transfection – PC3: 20 nM docetaxel or 18nM cabazitaxel for 72 hours; DU145: 8 nM docetaxel or 4 nM cabazitaxel for 48 hours. Cell viability was assessed with the CellTiter-Glo Luminescent Cell Viability Assay (Promega) at 1:2 dilution. The viability of cells transfected with microRNA mimic or inhibitor was represented by luminescence measurements normalised to those of cells transfected with the non-targeting control of respective drug treatments. MicroRNAs known to be associated with cardiotoxicity^57^ were eliminated from the dataset.

### Drug dose response curve

Reverse transfection of PC3 cells with microRNA mimics or non-targeting control was performed in 96-well microplates with 3000 cells and 0.08 μl of DharmaFECT-1 Transfection Reagent per well. At 24 hours post-transfection, culture medium was replaced with fresh medium. At 2 days post-transfection, cells were treated with a range of docetaxel (2-fold increments up to 150 nM) or cabazitaxel (2-fold increments up to 20 nM) doses for 48 or 72 hours respectively. Cell viability was assessed with alamarBlue Cell Viability assay (Invitrogen, ThermoFisher Scientific) by fluorescence detection according to the manufacturer’s instructions.

### Flow cytometry analysis of apoptosis and cell cycle

Labelling of apoptotic cells was performed using the Annexin V-FITC Apoptosis Detection Kit (BioVisionResearch Products, USA) according to manufacturer’s instructions. Cells were prepared for cell cycle analysis by fixing in 70% ethanol at −20^°^C, followed by staining with 0.1% propidium iodide (Sigma-Aldrich) and 830μg/ml RNase A (Sigma-Aldrich). Flow cytometry analysis was performed on the BD FACS Canto II system (BD Biosciences) using the FACSDiva software version 8. Data was analysed with FlowJo version 7.6.5.

### RNA sequencing and analysis

Total RNA was extracted from PC3 cells transfected with microRNA mimics or non-targeting control (4 replicates per treatment) using the Qiagen RNeasy mini kit (Qiagen, Germany) according to manufacturer’s instructions. Sequencing libraries were prepared with 1μg of intact RNA using the TruSeq Stranded mRNA Sample Preparation kit (Illumina) according to the manufacturer’s protocol. Sequencing was performed on the Illumina Hiseq2500 (125bp paired-end reads), with 12 samples multiplexed in one lane at approximately 20 million paired-end reads per sample. Raw reads were firstly assessed by FastQC and FastQ screen, then filtered using FastQ-mct (Github repository). Filtered reads were aligned to the human reference genome GRCh38 using STAR software version 2.4.2^58^. Feature count was obtained using RSEM version 1.2.21qq^59^.

Transcripts with significantly different levels between cells transfected with microRNA mimic and those with non-targeting control were identified using EdgeR package version 3.14.0^60^. Significantly different transcripts with extremely low RPKM (reads per kilobase per million) for both non-targeting and mimic-transfected cells (RPKM < 1), and low variance among the samples (variance < median variance of non-targeting and mimic-transfected cells) were removed before analysis for significant biological themes. Enriched gene sets were identifed using the Molecular Signatures Database (Broad Institute, Harvard, USA)^21^.

### siRNA knockdown

High-throughput siRNA knockdown of 51 different genes in PC3 cells was performed in 384 well microplates using the Caliper Zephyr Liquid Handling Workstation (Caliper Life Sciences, USA) and BioTek EL406 microplate dispenser/washer (BioTek Instruments, USA). All siRNAs were from the ON-TARGETplus SMARTpool collection (GE Healthcare Dharmacon). Reverse transfection of PC3 with the siRNAs was performed in the same conditions as for the microRNA screen, with four replicate wells per treatment. At two days post-transfection, culture medium was replaced with fresh medium containing 10 nM docetaxel, 5 nM cabazitaxel or drug-free medium for 72 hours. The miR-217 and miR-181b-5p mimics were included as positive controls. Cell viability was assessed with the CellTiter-Glo Luminescent Cell Viability Assay (Promega, Australia) as for the microRNA screen.

### Analysis of gene targets and public datasets

Prediction of microRNA gene targets was performed using our database (http://microrna.garvan.unsw.edu.au/mtp/search/index)^19^, which searches and collates the results from five other target prediction databases – StarBase version 2.0, miRTarBase version 4.5, TargetScan version 6.2, miRDB (accessed on April 2013), and DIANA microT-CDS version 5.

Microarray data of mRNA or microRNA expression were obtained from the cBioPortal database^26^ or NCBI GEO repository. Genes with more than 50% missing values in a tissue set were removed. Missing values were replaced with the minimum value of the dataset. Multiple t-tests was performed in Microsoft Excel (version 14.7.3), with Benjamini-Hochberg adjustment of P-values.

## Acknowledgements

This study was funded by Prostate Cancer Foundation of Australia (NCG-4212 and Movember Revolutionary Team Award MRTA1), and the Australian Prostate Cancer Research Centre-NSW through funds from the Australian Department of Health. The Victorian Centre for Functional Genomics (K.J.S.) was funded by the Australian Cancer Research Foundation (ACRF), the Victorian Department of Industry, Innovation and Regional Development (DIIRD), the Australian Phenomics Network (APN), and supported by funding from the Australian Government’s Education Investment Fund through the Super Science Initiative, the Australasian Genomics Technologies Association (AGTA), the Brockhoff Foundation and the Peter MacCallum Cancer Centre Foundation.

## Author information

**Contributions:** HML designed and performed experiments, analysed data, and wrote the manuscript; IN designed and performed experiments; JY, LC, CLC, NKY, CS and ZB performed experiments; ND analysed data; BE analysed data and developed database; ED and KJS designed protocols; BYL and RJD contributed data; LGH supervised the study; AS designed and supervised the study.

